# Gut Microbiota Alterations and Probiotic Intervention in Asian Elephants (*Elephas maximus*) with Gastrointestinal Distress

**DOI:** 10.64898/2026.06.22.733804

**Authors:** Ana Gabriela Herrera, Jacob Pederson, Stephanie Nuss, Vini Karumuru, Sehajvir Singh, Andrey Morgun, Richard R. Sim, Carlos R. Sanchez, Natalia Shulzhenko

## Abstract

Advances in metagenomic sequencing have transformed our understanding of host-associated microbiota, revealing critical roles in immune regulation, nutrient metabolism, and gastrointestinal (GI) homeostasis. However, the gut microbiome of large non-model species, particularly elephants, remains poorly characterized. Captivity introduces dietary, environmental, and management-related perturbations that may disrupt microbial balance and contribute to GI dysfunction.

Here, we performed a longitudinal analysis of the fecal microbiome in five captive Asian elephants (*Elephas maximus*) at the Oregon Zoo exhibiting chronic fecal abnormalities, including mucus and inconsistent stool formation. Over 14 weeks, weekly fecal samples were collected and compared with samples from clinically normal elephants housed at three other zoological institutions. Using 16S rRNA gene sequencing, we identified marked differences in microbial community composition between affected and control elephants. Dysbiosis in Oregon Zoo elephants was characterized by enrichment of *Akkermansia muciniphila* and multiple members of the order *Clostridiales*, including taxa previously associated with gastrointestinal disorders.

Administration of a commercially available probiotic formulation was associated with transient improvement in fecal characteristics and pronounced shifts in microbial composition, including a significant post-treatment reduction in overall microbial diversity and decreased abundance of several taxa linked to GI abnormalities. Notably, probiotic strains themselves were not detected, suggesting indirect or short-lived functional effects rather than durable colonization.

Together, these findings provide one of the first longitudinal characterizations of gut microbiome dysbiosis in captive Asian elephants and identify candidate microbial contributors to chronic GI dysfunction in captivity, with implications for husbandry, dietary management, and microbiome-informed interventions in megafauna. Additionally, our study underscores the potential, although limited and likely indirect, benefit of probiotics when treating GI disorders in monograstric megavertebrates.

## Introduction

The gut microbiome—comprising bacteria, archaea, viruses, protozoa, and fungi—forms a complex and dynamic ecosystem that profoundly influences host physiology^[1]^. Beyond its central role in digestion, the gastrointestinal (GI) microbiota contributes to immune development, energy metabolism, epithelial barrier integrity, and neuroendocrine signaling^[2,3]^. Microbial activities such as short-chain fatty acid production, vitamin synthesis, pathogen exclusion, and modulation of host immune responses make the microbiome an essential component of host^[4–6]^.

While human and laboratory animal microbiome research has expanded rapidly, comparatively little is known about the gut microbiota of large non-model mammals, particularly megaherbivores such as elephants^[7–10]^. Captivity imposes substantial alterations in diet diversity, environmental microbial exposure, stress, and management practices, all of which can reshape gut microbial communities^[11]^. Gastrointestinal dysfunction is among the most common chronic health concerns in zoo-housed elephants and frequently manifests as persistent diarrhea, abnormal stool consistency, or excessive fecal mucus^[12]^.

Asian elephants (*Elephas maximus*) are obligate hindgut fermenters that depend heavily on a complex colonic microbial community for the degradation of fibrous plant material. Diet composition and plant diversity are therefore critical determinants of gut microbial structure and function. A recent study demonstrated that plant diversity in elephant diets decreases with increasing degrees of captivity, potentially predisposing animals to microbiome instability. Disruption of these microbial communities may compromise mucosal integrity and fermentation efficiency, contributing to chronic GI signs and can manifest in other health effects^[13,14]^.

The present study aimed to (i) characterize the fecal microbiome of Asian elephants under human care exhibiting chronic fecal abnormalities, (ii) compare these microbial communities with those of clinically normal elephants housed at other institutions, and (iii) assess longitudinal microbiome and clinical responses to probiotic administration. By integrating microbial profiling with detailed clinical observations, this work seeks to advance understanding of microbiome-associated GI dysfunction in captive elephants.

## Methods

### Animals and Clinical Background

Five adult Asian elephants (*Elephas maximus*), 2 males and 3 females housed at the Oregon Zoo (ORZ), Portland, Oregon, were included in this study (Figure 1B). Beginning in September 2020, animal care staff observed intermittent fecal abnormalities, including excessive mucus and inconsistent bolus formation, which persisted cyclically for approximately two years. Comprehensive clinical evaluations revealed no evidence of systemic illness: hematology and serum biochemistry values were within reference ranges, and repeated fecal cytology and fecal analysis including aerobic, anaerobic and fungal culture and polymerase chain reaction testing that showed no evidence of inflammatory, bacterial, fungal, or parasitic disease. The protozoan *Trophozoite* was detected in rare to moderate levels in some samples but was not associated with pathological findings and it is considered a normal protozoon inhabiting the caecum and colon of Asian and African elephants^[15]^.

**Figure 1.**
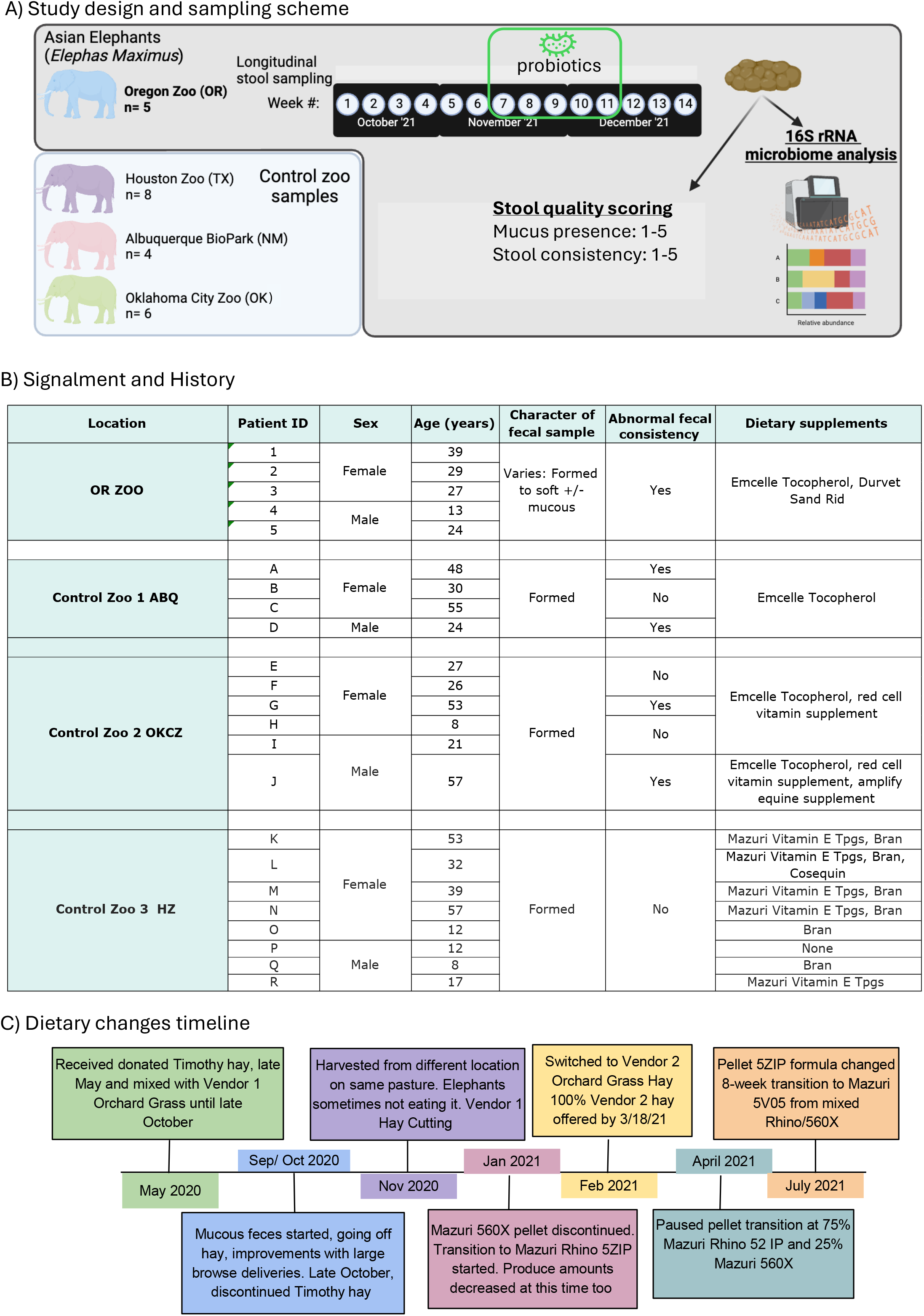
(A) Study design and sampling scheme from Oregon Zoo (ORZ) elephants (gray box) and control zoo elephants (blue box). In ORZ elephants, stool samples were collected weekly over the course of 14 weeks. Probiotics were administered weeks 7-11. Control stool samples (1 per elephant) were collected from 18 elephants across 3 other zoos. All stool samples underwent quality scoring (mucus presence, stool consistency) and 16S rRNA sequencing, followed by microbiome analysis. (B) Table of signalment and history of elephants, separated by zoo. (C) Timeline of dietary changes in ORZ elephants, detailing transitions in hay and pellet type/origin.

Given the absence of identifiable pathogens, the lack of systemic clinical signs or bloodwork abnormalities, a non-infectious etiology involving microbial imbalance was suspected. Fecal glucocorticoid (fG) metabolite measurements were compared with baseline values to determine whether stress could be associated to the fecal abnormalities, but fG remained within baseline ranges. Feed quality and nutritional composition were evaluated repeatedly and met established dietary standards.

### Probiotic Administration

To try to mitigate persistent mucoid feces and support microbial balance, oral probiotic therapy was initiated during week 7 of the study (November 2021). All elephants received 60 grams ProBios® Bovine Oral Gel (Vets Plus Inc., Menomonie, WI), containing *Enterococcus faecium, Lactobacillus acidophilus, L. casei*, and *L. plantarum* (≥10^7^ CFU/g), administered once a day for 28 consecutive days (four consecutive weeks (weeks 7–10)). The same formulation and dosing schedule were used for all individuals.

### Diet and Management

Diet composition and feeding records were monitored throughout the study by the zoo’s nutritionist. Routine inspections were conducted at feed delivery, storage, and prior to feeding. Several changes in hay supplier and manufacturer occurred prior to the onset of GI signs (Fig. 1C). Although nutritional analyses indicated compliance with dietary standards, detailed records were not consistently available.

### Sample Collection

Fecal samples were collected weekly in the same manner for 14 weeks, from October 2021 to January 2022 (Fig. 1B). A total of 88 samples were analyzed: 70 samples from ORZ elephants (5 individuals × 14 weeks). Control samples were collected from clinically normal Asian elephants housed at the following institutions: Houston Zoo (HZ) in Texas state, Albuquerque BioPark (ABP) in New Mexico state, and Oklahoma City Zoo (OKCZ) in Oklahoma state. Samples were collected immediately after defecation and stored at −80 °C.

Fecal consistency was scored on a five-point scale (1 = dry, 5 = liquid), and mucus presence was assessed using a standardized visual scoring system. Scoring was performed consistently by trained personnel. Control institutions reported no recent GI abnormalities. Signalment and medical histories are summarized in Figure 1B.

### DNA Extraction and Sequencing

DNA was extracted using the Qiagen PowerFecal DNA Stool Kit according to the manufacturer’s protocol. DNA concentration was quantified using a Qubit 2.0 fluorometer (Thermo Fisher Scientific). The V4 hypervariable region of the 16S rRNA gene was amplified using primers 515F and 806R, with unique barcodes assigned to each sample. Amplicons were visualized by agarose gel electrophoresis, purified, and sequenced at the Center for Quantitative Life Sciences, Oregon State University, using an Illumina MiSeq platform with 250 bp paired-end reads.

### Bioinformatic and Statistical Analysis

Sequencing data were processed using the DADA2 pipeline implemented in QIIME2 (v2023.2) to infer amplicon sequence variants (ASVs)^[16,17]^. Taxonomic classification was performed using the SILVA 138 reference database^[18]^. Alpha diversity was assessed using the Chao1 index, and beta diversity was calculated using Bray–Curtis dissimilarity^[19]^.

Differential abundance analysis was performed using DESeq2 with a false discovery rate (FDR) threshold of 0.2, consistent with standard practice for exploratory microbiome studies with small sample sizes^[20]^.Longitudinal associations between microbial taxa and fecal traits were evaluated using MaAsLin2, accounting for repeated measures within individuals and relevant covariates^[21]^.

## Results

Eighty-eight fecal samples were analyzed, including 70 samples from ORZ elephants and 18 samples from control elephants (Fig. 1A). ORZ elephants exhibited intermittent mucus production and inconsistent stool formation throughout the study period but there were no dietary changes during the period of sample collection (Fig. 1C).

Alpha diversity, assessed using the Chao1 index, increased during the probiotic administration period (weeks 8–11) and declined significantly following cessation of treatment (Fig. 2A).

**Figure 2.**
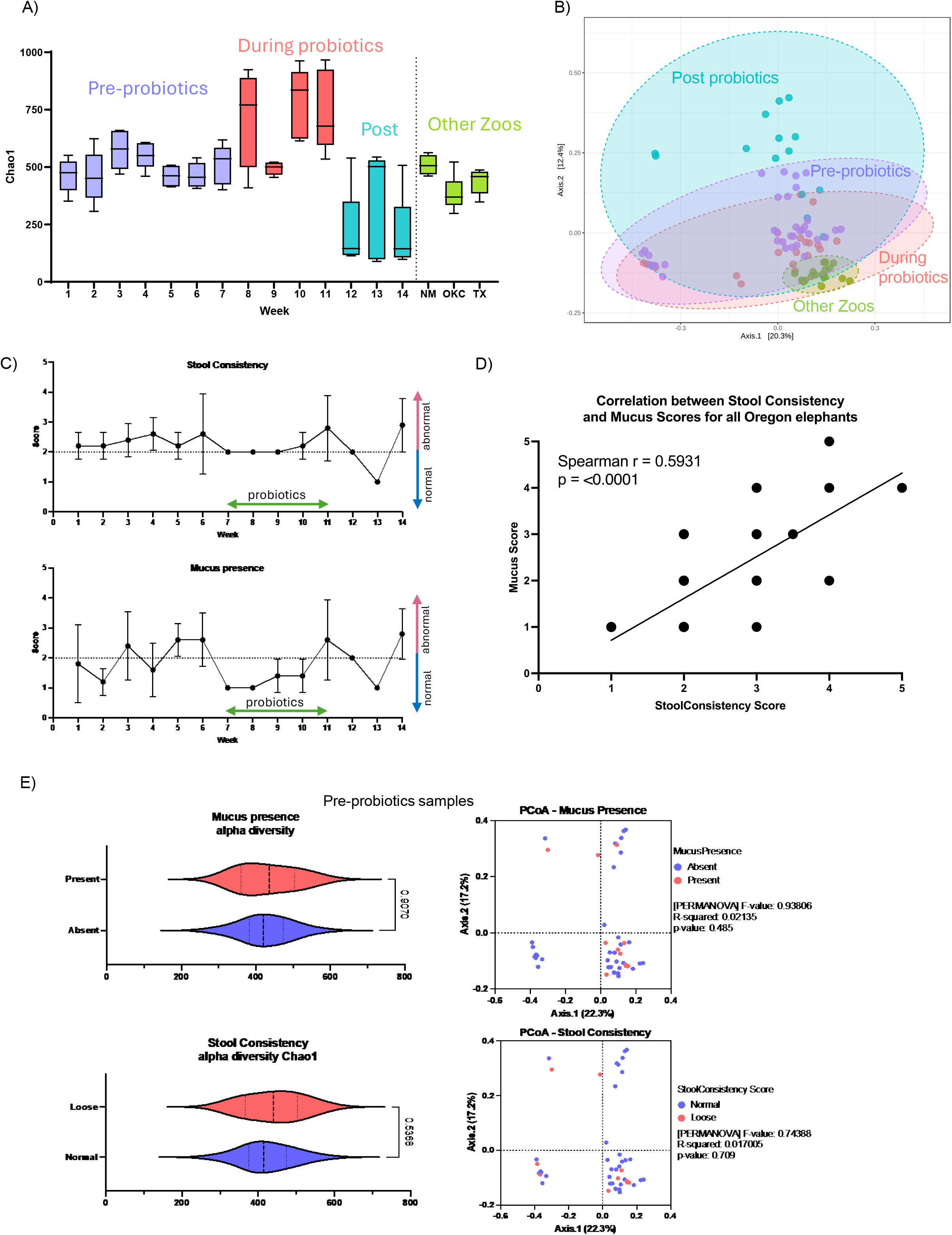
(A) Distribution of Chao1 diversity index (alpha diversity) at each week in ORZ elephants (left of dotted line) versus control elephants from other zoos (green; right of dotted line). Boxes corresponding to ORZ elephants are colored in relation to probiotic administration. (B) PCoA plot based on Bray-Curtis dissimilarity (beta diversity) in ORZ elephants (purple, red, blue) and control elephants (green). (C) Stool consistency and mucus presence scores of ORZ elephants over time. (D) Mean mucus presence score vs. mean stool consistency score of ORZ elephants. Each point represents a week. (E) Alpha diversity (Chao1 index) and beta diversity (Bray-Curtis dissimilarity) measures of normal (blue) vs. abnormal (red) in pre-probiotic (weeks 1-6) ORZ stool samples. Normal samples have scores ≤ 2 (no mucus or normal stool consistency), whereas abnormal samples have scores >2 (presence of mucus or loose stool consistency). Statistics shown correspond to student’s t-tests (for Chao1 index) and PERMANOVA tests (for Bray-Curtis dissimilarity) to compare normal vs. abnormal samples.

Beta-diversity analysis using Bray–Curtis dissimilarity revealed clear separation between ORZ elephants and controls, independent of probiotic treatment phase, whereas samples from control institutions clustered together (Fig. 2B).

Temporal trends showed parallel improvement in fecal consistency and mucus scores during probiotic administration (Fig. 2C), with a significant correlation between these two clinical measures (Fig. 2D). However, within ORZ elephants, beta-diversity did not differ significantly between samples classified as clinically normal or abnormal (Fig. 2E), suggesting persistent underlying dysbiosis.

To identify candidate taxa associated with GI dysfunction, we compared samples collected before the use of probiotics with those from the other zoos to exclude this potential confounding (Fig. 2A). Among 233 taxa present at ≥0.1% relative abundance, 89 taxa (38%) differed significantly between groups (FDR < 0.2), with 64 taxa enriched and 25 taxa depleted in ORZ elephants (Fig. 3A).

**Figure 3.**
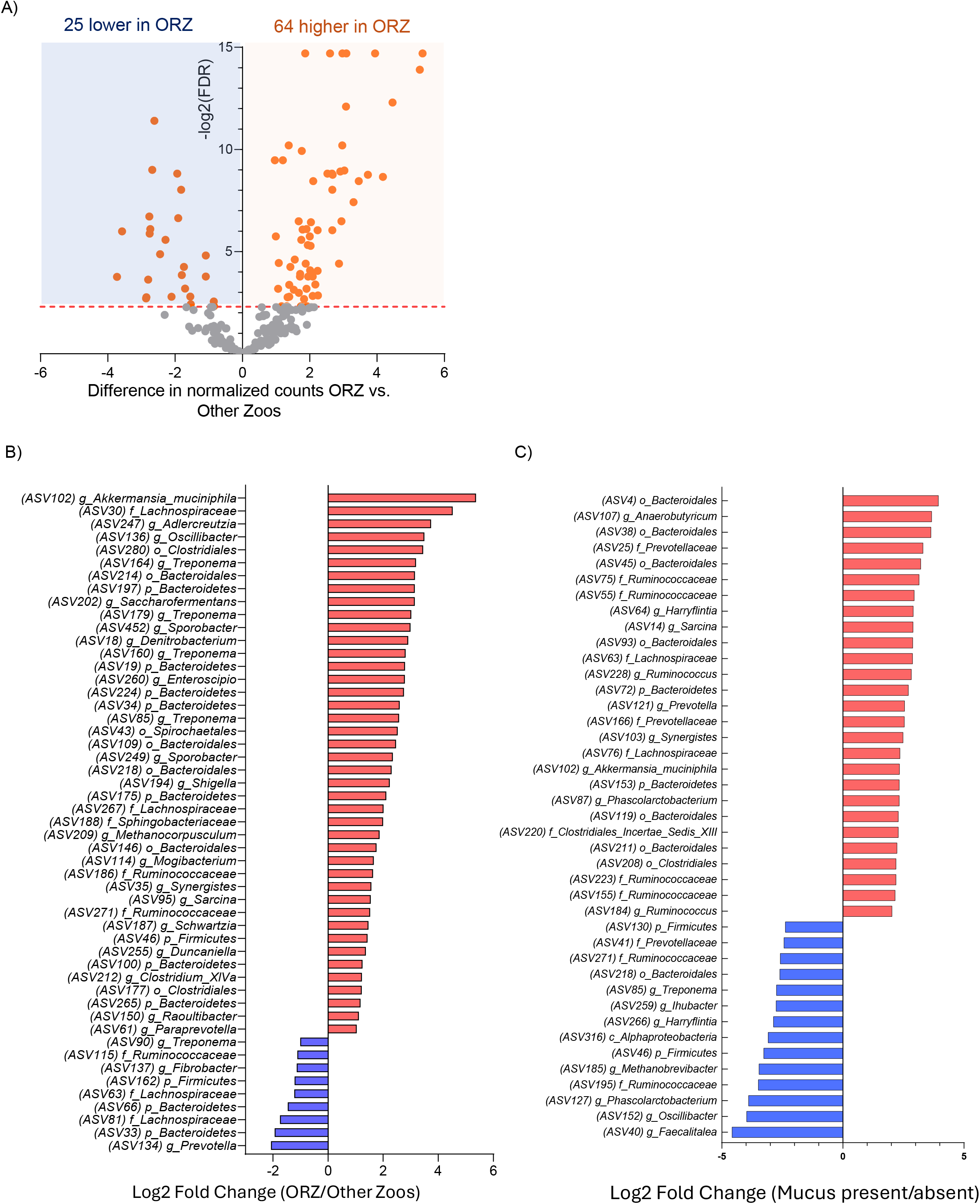
(A) Volcano plot showing difference in mean normalized counts per taxa in ORZ elephants (pre-probiotic samples) versus control elephants. Taxa present at ≥0.1% relative abundance are shown, with taxa that have FDR < 0.2 colored orange. Colored boxes distinguish direction of fold change (light blue: lower abundance in ORZ elephants; light orange: higher abundance in ORZ elephants). (B) Log_2_ fold change of most enriched taxa in pre-probiotic stool samples from ORZ elephants (highest magnitude of fold change relative to elephants from other zoos). (C) Log_2_ fold change (comparing stool samples with versus without mucus) of 56 taxa, identified with MaAsLin2, that were significantly associated with mucus presence in ORZ elephants after adjusting for elephant identity, stool consistency, and sampling week.

Notably, *Akkermansia muciniphila* (a mucin-degrading bacterium linked to GI mucosal thinning and low-fiber diets) was among the most enriched taxa in ORZ elephants (Fig. 3B)^[22]^. Several other enriched taxa belonged to the order *Clostridiales*, which includes species previously implicated in GI disorders^[23]^.

MaAsLin2 analysis of pre-probiotic ORZ samples identified 56 ASVs significantly associated with mucus presence after adjusting for elephant identity, stool consistency, and sampling week (Fig. 3C). No ASVs were significantly associated with stool consistency after accounting for these covariates.

To increase robustness in this longitudinal dataset, we applied an averaging-across-timepoints approach previously used in human inflammatory bowel syndrome studies^[24]^. For each elephant, samples from weeks 1–7 were classified as either “abnormal” or “normal,” then taxonomic abundances were averaged within each group and then compared in a pairwise manner, for each elephant. This analysis identified 29 differentially abundant taxa, of which 9 were increased in abnormal samples (Fig. 4A).

**Figure 4.**
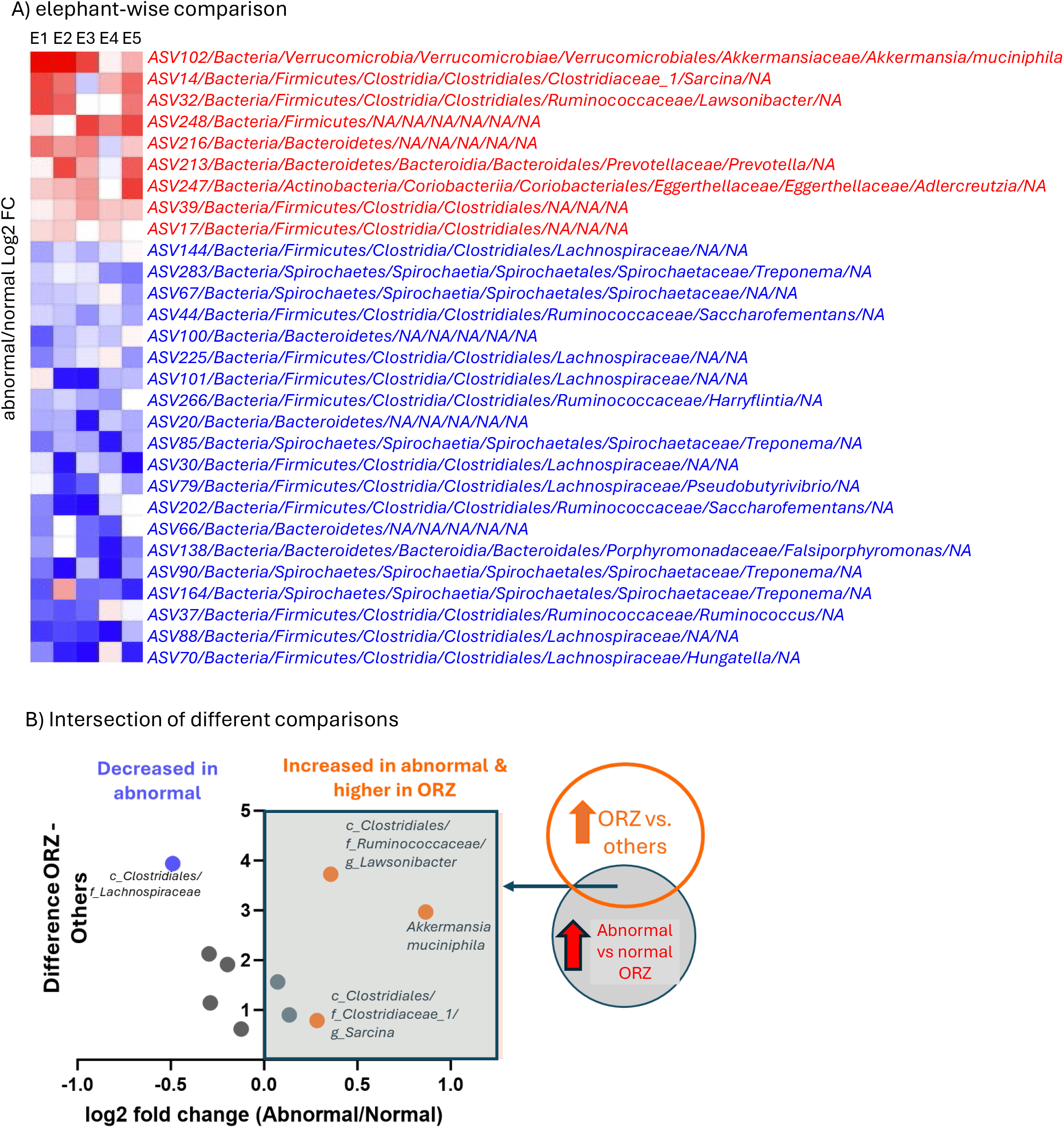
(A) Heatmap of pairwise comparisons between normal and abnormal samples within each Oregon Zoo elephant. 29 identified differentially abundant taxa are shown. Red taxa are more abundant in abnormal samples (log_2_FC > 0), while blue taxa are more abundant in normal samples (log_2_FC < 0). (B) Intersection of taxa with those differentially abundant in ORZ elephants relative to controls with five ASVs with >0.5% abundance that were enriched in ORZ.

Intersection of these taxa with those differentially abundant in ORZ elephants relative to controls yielded five ASVs with >0.5% abundance that were enriched in ORZ (Fig. 4B). *Akkermansia muciniphila* was again the most prominent taxon, while the remaining four ASVs belonged to the order *Clostridiales*, including species of *Sarcina* reported as a rare opportunistic pathogen^[25]^.

Comparison of average taxonomic abundance between pre- and post-probiotic periods identified 36 taxa with significant changes (Fig. 5). All but one taxon (*Treponema*, ASV1) decreased following probiotic administration. Importantly, *Akkermansia muciniphila* was among the taxa showing the greatest decline in the post-probiotic period. Taxa corresponding to species contained in the probiotic formulation were not detected, indicating that clinical and microbial effects were likely indirect.

**Figure 5.**
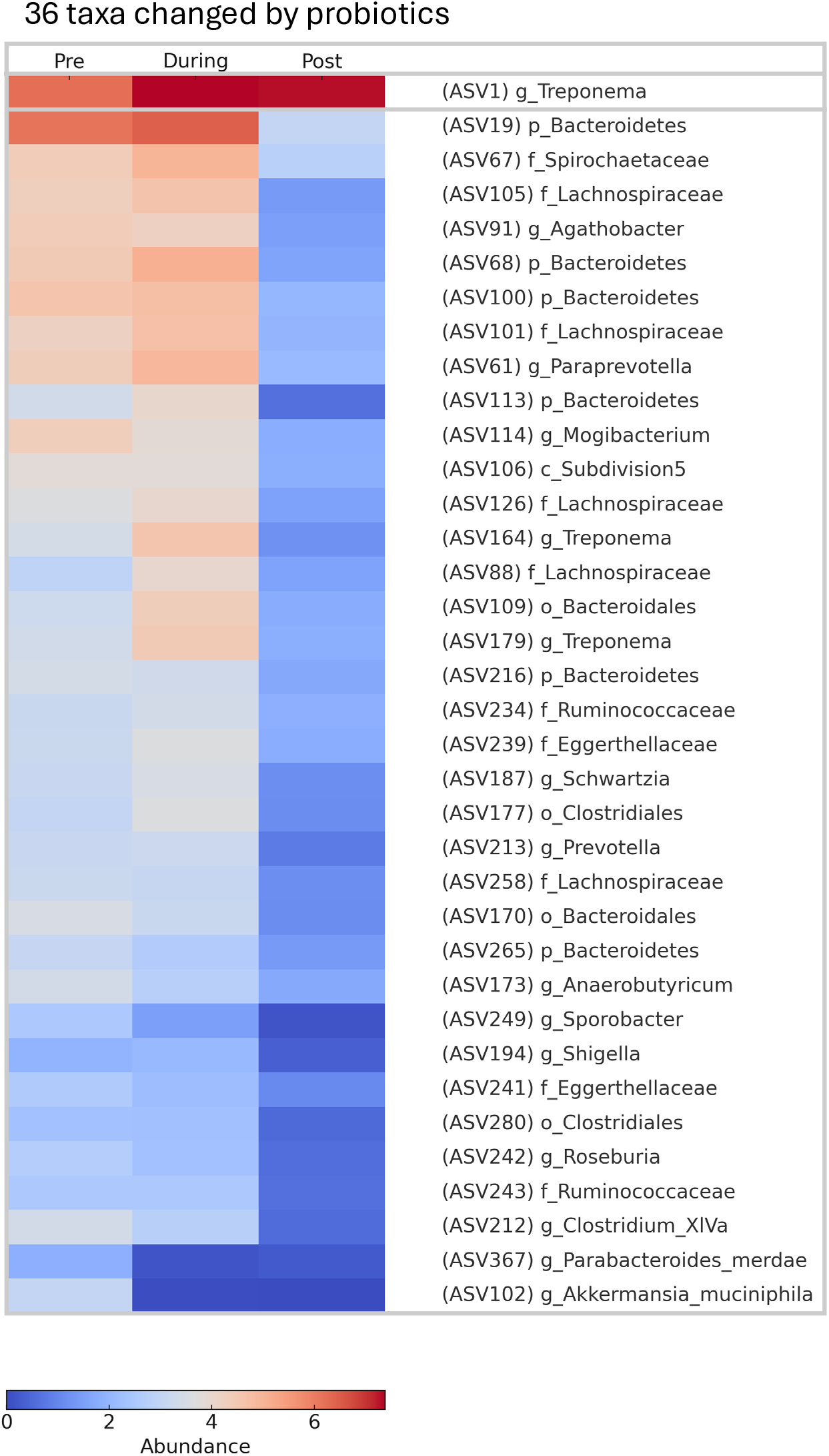
Heatmap of 36 taxa whose mean abundances changed between pre- and post-probiotic periods in ORZ elephants.

## Discussion

This study provides one of the first longitudinal characterizations of the gut microbiome in captive Asian elephants exhibiting chronic GI abnormalities. Our findings reveal persistent dysbiosis in affected elephants, transient clinical and microbial responses to probiotic administration, and consistent enrichment of candidate taxa—particularly *Akkermansia muciniphila* and members of *Clostridiales*—that may contribute to GI pathology.

A striking observation was the non-monotonic response of alpha diversity, which increased during probiotic administration but declined markedly after treatment cessation. While probiotics are often expected to enhance microbial diversity, the administered formulation consisted primarily of lactic acid-producing bacteria that are not typical members of the elephant hindgut microbiome. The transient improvement in fecal quality during treatment, coupled with the absence of detectable probiotic strains, suggests short-term functional modulation rather than stable microbial engraftment.

Dietary and environmental factors likely contributed to the observed dysbiosis such as changes in hay suppliers, which were in part related to the wildfires in Oregon in early September 2020, a period that preceded the GI sign onset in elephants. Although nutritional analyses met established standards, incomplete historical records of browse composition limit retrospective evaluation of plant diversity and seasonal variation. In other herbivores and livestock species, subtle feed quality differences are known to drive microbial shifts favoring opportunistic taxa^[26]^.

Across multiple analyses, *Akkermansia muciniphila* showed the strongest and most consistent association with fecal mucus. This mucin-degrading bacterium has been linked to colonic barrier thinning under low-fiber conditions and may be particularly disruptive in hindgut fermenters that depend on an intact mucus layer for fermentation efficiency^[22]^. Supporting this interpretation, recent work in elephants identified *Akkermansia* exclusively in diarrheic animals but not in those with colic, suggesting a specific association with mucosal disruption rather than motility disorders^[27]^.

In addition, several *Clostridiales* taxa, including those related to *Sarcina*, were consistently enriched in abnormal samples. *Sarcina* species are spore-forming bacteria capable of surviving harsh conditions and producing microbial cellulose, which may facilitate *Akkermansia* expansion. Although their pathogenicity in elephants is unknown, *Sarcina* has been associated with severe GI disease in other animals supporting its candidacy as a contributing factor^[25,28,29]^.

Together, these observations support a working model in which dietary or environmental perturbations introduce or favor opportunistic taxa that can stimulate mucus production as a host defense. This altered mucosal environment may then promote expansion of mucin-degrading bacteria, creating a feedback loop that perpetuates barrier dysfunction and chronic GI abnormalities.

Study limitations include the small cohort size, geographic and diet differences between ORZ and control institutions, and the absence of contemporaneous microbiome data from elephants after clinical recovery. Future work incorporating metagenomic sequencing, dietary intervention studies, and longitudinal follow-up after GI sign resolution will be essential to validate proposed mechanisms.

## Conclusion

Our findings suggest that environmental and dietary perturbations may initiate a cascade of microbial imbalances that contribute to chronic GI dysfunction in captive Asian elephants. The identification of specific taxa associated with fecal mucus and abnormal stool highlights potential microbial targets for diagnostics and intervention. Although probiotic administration produced transient clinical improvement and substantial microbial shifts, these effects did not reflect durable restoration of a stable hindgut microbiome. These results underscore the importance of consistent diets, detailed dietary record keeping and microbiome-informed management strategies for the health of captive megafauna.

## Data availability

Raw data have been deposited to the NCBI Sequence Read Archive under BioProject accession number PRJNA1480372.

## Acknowledgements

The authors would like to sincerely thank the following individuals and institutions for their contribution to this study: Carol Bradford (Albuquerque BioPark), Christine Molter (Houston Zoo), Jennifer D’Agostino & Gretchen Cole (Oklahoma City Zoo), and the elephant staff and veterinary staff at the Oregon Zoo for sample collection, processing and shipment.

